# P2X7 receptor-mediated astrocytic atrophy in the hippocampus of mice after *status epilepticus*

**DOI:** 10.64898/2026.04.16.718853

**Authors:** Xuan Li, Muhammad Tahir Khan, Elek Sylvester Vizi, Beata Sperlagh, Si-Si Lin, Alexej Verkhratsky, Patrizia Rubini, Yong Tang, Peter Illes

## Abstract

Genetic deletion or pharmacological blockade of P2X7 receptors (Rs) counteract *status epilepticus* (SE) in animal models of epilepsy. It is, however, unclear whether P2X7Rs are localized at astrocytes or neurons, and the reason for astrocytic atrophy arising in consequence of SE is also ambiguous. We conducted a combined morphological/electrophysiological study in order to investigate these issues. It has been shown that kainic acid (KA)-induced SE in mice led to the atrophy of hippocampal astrocytes and at the same time to the decrease of ezrin immunoreactivity and its co-expression with mCherry, whose synthesis has been initiated by the injection of a virus complex. mCherry expression in astrocytes enabled us to study changes in cell somata and processes brought about by KA-injection. Ezrin is a plasmalemmal-cytoskeleton linker; its grade of expression indicates changes in the existence/function of small peripheral astrocytic processes. Pretreatment of mice with the blood-brain barrier-permeable P2X7R antagonist JNJ-47965567 prevented the SE-induced damage of astrocytes. KA caused a potentiation of dibenzoyl-ATP (Bz-ATP) currents in astrocytes but not neurons of the hippocampus. This effect was also abolished by pre-treatment of mice with JNJ-47965567 before applying KA, although no similar changes occurred in hippocampal CA1 neurons. The measurement of spontaneous postsynaptic currents (sPSCs) and spontaneous excitatory postsynaptic currents (sEPSCs) indicated a presynaptic facilitation of neurotransmitter release by Bz-ATP. In conclusion, we suggest that astrocytic P2X7Rs are the primary target of ATP release from damaged CNS cells in the hippocampus which simultaneously causes damage to astrocytic somata and processes.

## Introduction

Epilepsy is characterized by recurrent spontaneous seizures, caused by hyperexcitability of cerebro-cortical neurons, generally believed to be due to an imbalance of synaptic excitation and inhibition [1, 2]. Although epilepsy is a common neurological disease broadly affecting the general human population with an incidence of 1-2% and results in significant limitations in personal and social activities, its extreme form, *status epilepticus* (SE), is associated with considerable morbidity and mortality [3, 4]. Continuous grand mal seizures lasting longer than 5 min, or absence/focal epilepsy continuing for >20-30 min without recovering consciousness is defined as SE [5].

Epilepsy has been considered a neuronal disease, but more recently glial cells, especially astrocytes and microglia are considered to be important factors in initiating recurrent seizures *via* altering neuronal homeostasis [6–8]. Astrocytes represent 20-40% of the cells in the brain, serving a multitude of functions, most notably ion homeostasis, neurotransmitter clearance, lipid homeostasis, synapse formation/removal, synaptic modulation and contribution to neurovascular coupling [9, 10]. Protoplasmic astrocytes located in the cerebro-cortical grey matter relevant for the regulation of epilepsy, possess a cell body, six or more major branches that emanate from the soma, and further fine branches and leaflets that contact synapses usually called perisynaptic or peripheral astrocyte processes (PAPs) [10, 11]. These PAPs are below the resolving power of light microscopy but are especially important for constituting the contact with neuronal synapses and form the so-called synaptic cradle. The morphological plasticity of leaflets is regulated by the plasmalemmal-cytoskeleton linker ezrin, involved in astrocyte motility [12, 13].

In 1972, Geoffrey Burnstock proposed the existence of ‘purinergic signaling’ as a way of intercellular communication [14]. It had been demonstrated that adenosine 5’-triphosphate (ATP) and its metabolite adenosine exert concentration-dependent inhibitory effects in the nerve-stimulated guinea-pig myenteric plexus-longitudinal muscle preparation [15]. According to the hypothesis of Burnstock [16], the effect observed on both smooth muscle and nerve cells was thought to be mediated by P1 (adenosine) and P2 (ATP) receptors. Subsequently, twenty-one distinct P2 receptors were identified, and it was shown that ATP, whether released via synaptic vesicles or from the postsynaptic membrane, plays an active role in neurotransmission [17–19].

In conclusion, the effects of ATP have been shown to be mediated by two types of purinoceptors (P2X, ligand-gated cation channels; P2Y, G-protein-coupled receptors), and modified by the release of ATP from neurons/astrocytes, and the degradation of ATP by metabolizing enzymes [16, 20, 21]. It is especially important to note, that ATP is a glio-transmitter released by exocytotic and non-exocytotic mechanisms from astrocytes and thereby establishes the ‘tripartite synapse’, consisting of the presynaptic neuronal terminals, the postsynaptic neuronal specializations and small astrocytic processes [11, 22, 23].

Of the P2X receptor (R) family, the P2X7R subtype has been reported to be intimately associated with the pathophysiology of SE [24–26]. Extensive evidence supports the assumption that these findings might have broad therapeutic consequences; P2X7R pharmacological antagonists acted beneficially not only in animal models of epilepsy, but were also shown to be possible adjunct therapeutic agents in drug-resistant epilepsy [25, 27], had promising qualities as diagnostic biomarkers for temporal lobe epilepsy [28], and were shown even to prevent co-morbidities of epilepsy such as cognitive deterioration [29, 30].

Neuroinflammation [31, 32] and astroglial scarrring [33] are causative factors to induce epilepsy and SE. In this context it perfectly fits that P2X7R activation by massively released ATP causes neuroinflammation/gliosis [34, 35], and the blockade or genetic deletion of P2X7Rs has anti-epileptic effects (see above). It has been reported that SE in rodent models is associated with atrophy of distal astrocytic branches and reduced astrocyte coupling in their syncytium [36, 37]. Based on these findings we asked ourselves, whether kainic acid (KA)-induced SE in mice leads to astrocytic atrophy and consequent impairment of astrocytic function *via* the activation of P2X7Rs by the release of large amounts of ATP and whether astrocytic ezrin is involved in this process. All questions could be answered in an affirmative manner.

## Materials and Methods

### Animals

C57BL/6J male mice (from Chengdu Dossy Experimental Animal Co., Chengdu, China), 7 weeks old, were used for the experiments reported in this paper. All mice were adapted to the standard laboratory conditions, which were 24±2°C room temperature and 65±5% humidity on 12/12h light/dark cycles, with drinking water and food available *ad libitum*. The animal study was reviewed and approved by the Institutional Review Board of the Chengdu University of Traditional Medicine, Chengdu, China (protocol code, AF2475, 2020 April 1), and was in accordance with the National Institute of Health Guidelines for the Care and Use of Laboratory Animals.

### Epilepsy model

KA (30 mg/kg) was injected intraperitoneally (i.p.; Fig. 1). The dose of KA and its route of application were the same in all experiments included in this paper. Saline (0.9% NaCl) was injected to the control group instead of KA. In previous experiments a modified Racine scale was used for measurement of the intensity of seizures; this proved the efficiency of KA at the present dosage to induce SE [30]. Seizures at stages 4-5 of the Racine scale that last for ≥30 min were defined as SE.

**Figure 1.**
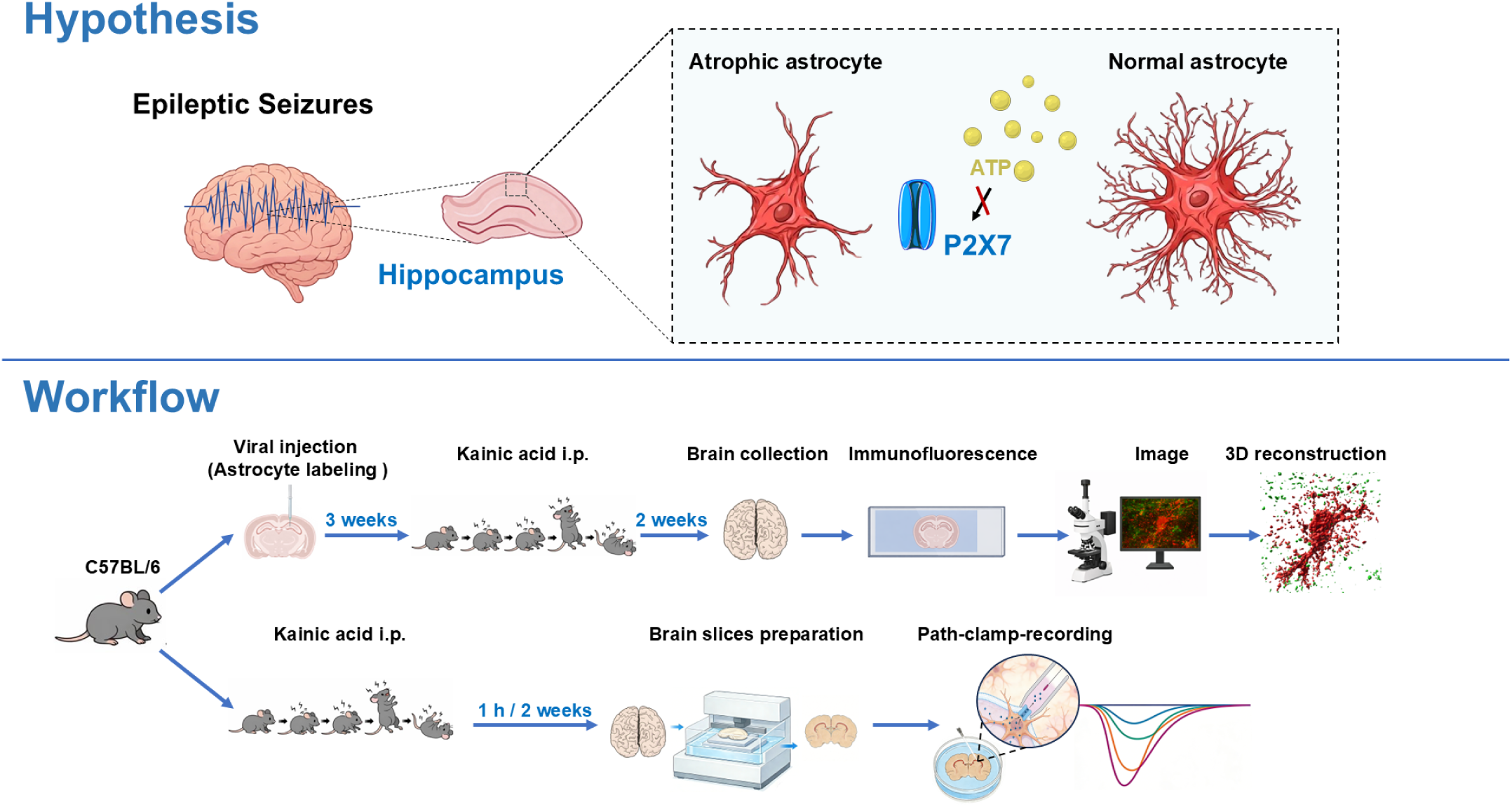
Experimental hypothesis and workflow for investigating astrocytic P2X7R receptor signaling in kainic acid–induced epilepsy. Hypothesis: Epileptic seizures trigger ATP release and P2X7 receptor activation, leading to morphological atrophy and functional impairment of hippocampal astrocytes. Workflow: Adult C57BL/6 mice received into the hippocampus viral injections for astrocyte-specific labeling with mCherry. After 3 weeks, epilepsy was induced by intraperitoneal (i.p.) injection of kainic acid (KA). For morphological analyses, brains were collected 2 weeks after KA administration, followed by immunofluorescence staining, image acquisition, and 3D reconstruction of astrocytes. In parallel, for electrophysiological recordings, acute hippocampal slices were prepared at 1 h or 2 weeks after KA treatment and subjected to patch-clamp recordings to assess astrocytic functional properties.

The mortality of mice was 10-15% in our experiments with 30 mg/kg, i.p. KA, and thereby comparable to that reported by Augusto [38] (35 mg/kg KA, subcutaneously). The mortality could have been reduced by the injection of diazepam, 1 h after injection of KA, to terminate seizures (e.g. [39]), although of course death during the 1 h period is unavoidable even in this case. Moreover, treatment with the sedative/antiepileptic diazepam might have interfered with astrocytic atrophy, which is not compatible with the present experimental aims. We relinquished from the application of diazepam, because the mortality was relatively low and death within the 1-h period before diazepam injection could not have been avoided either.

### Stereotaxic surgery and virus application

Mice were anesthetized with isoflurane (4% induction; 1-1.5% maintenance; RWD Life Science, Shenzhen, China) and fixed on a stereotactic platform (RWD Life Science). The animals were placed on heating pads (37°C) during surgery to keep their body temperature stable. A 1 μl Hamilton syringe fixed with a beveled glass pipette was inserted into the right side of the hippocampal CA1 region. The coordinates used were: AP −2.0 mm, ML +1.5 mm (right), and DV −1.45 mm from the skull surface. A total of 0.8 μl of AAV5-gfaABC1D-mCherry (1 × 10^12^ gc/ml; Taitool Bioscience, Shanghai, China), was slowly applied. Glass pipettes were left in place for at least 5 min. After injection, animals were allowed to completely recover under a warming blanket and then returned to the home cage. An interval of 3 weeks was kept between virus application and the induction of the epilepsy model (Fig. 1).

### 3D reconstruction

The 3D reconstruction of astrocytes in the hippocampal *stratum oriens* was performed as described previously [40, 41]. The confocal imaging stacks were collected with a Z-step size of 0.25 μm under a confocal microscope (Olympus, Tokyo, Japan). Three-dimensional reconstructions were processed offline using Imaris 10.1.0 (Bitplane, South Windsor, CT, USA) as reported by others [42]. In brief, the astrocyte soma and processes were measured and reconstructed according to their own parameter. The processes’ diameter was measured as one-tenth of astrocyte soma. In addition, ezrin was measured as 1 mm in every group. The surface-to-surface co-localization was calculated by a specific plugin of Imaris [43].

### Sholl analysis

Sholl analysis was performed on astrocytes using Imaris 10.1.0, with the built-in Sholl analysis model after three-dimensional reconstruction. The soma was defined as the center and concentric spheres were created at 1 µm radial intervals. The number of intersections between astrocytic processes and each sphere was automatically quantified.

### Immunofluorescence staining

Mice were perfused through the ascending aorta with cold paraformaldehyde (PFA, 4%w/v in phosphate buffered saline (PBS)) after the i.p. injection of 1% sodium pentobarbital (0.4 ml). The brains were removed and postfixed in 4% paraformaldehyde overnight. Then, they were dehydrated with gradient (20-30%) sucrose in PBS at 4°C. Coronal 40 μm-thick sections were prepared and incubated in a cryostat (Leica CM1860; Leica Biosystem, Muttenz, Switzerland) at −80°C until use. The sections were afterwards incubated in a blocking solution containing 4% bovine serum albumin (Sigma-Aldrich, Shanghai, China) and 0.5% Triton X-100 (Solarbio, Beijing, China) for 2 h at room temperature. Subsequently, the sections were incubated with the primary antibody (rabbit anti-ezrin 1:100; Cell Signaling, Danvers, MA, USA; mouse anti-GFAP; 1:500, Cell Signaling; mouse anti-S100β; 1:200, Abcam, Cambridge, MA, USA) overnight at 4°C. After washing in PBS three times, slices were incubated for 2 h in saturation solution containing the relevant secondary antibody (goat anti-rabbit Alexa 488; Invitrogen, Carlsbad, CA, USA). After washing in PBS three times, and labelling in some cases the cell nucleus with DAPI (1:10,000; Abmole Bioscience, Houston, TX, USA), the coverslips were mounted on slides using anti-fade solution (Solarbio, Beijing, China). Image acquisition was performed using a confocal laser scanning microscope (Olympus IXplore SpinSR, Olympus, Tokyo, Japan).

For GFAP or S100β quantification, six regions of interest (ROIs; 400 x 400 pixels) in the *stratum oriens* and four ROIs (800 x 800 pixels) in the *stratum radiatum* were randomly selected per mouse. Images were acquired at a spatial resolution of 0.311 μm per pixel. Cell counts were normalized and expressed as cell density (cells/mm^2^).

### Patch-clamp recording and agonist-induced currents

C57BL mice, 7 weeks old, were utilized for these experiments. 1 h or 2 weeks after inducing SE with the i.p. application of KA (Fig. 1), mice were sacrificed with the i.p. injection of 1% sodium pentobarbital (0.4 ml; Sigma-Aldrich). The preparation of the hippocampal slices and patch-clamp procedures were as described previously [44]. After decapitation, the brain was placed into ice-cold, oxygenated (95% O_2_ + 5% CO_2_) artificial cerebrospinal fluid (aCSF) of the following composition (in mM): NaCl 126, KCl 2.5, CaCl_2_ 2.4, MgCl_2_ 1.3, NaH_2_PO_4_ 1.2, NaHCO_3_ 25, and glucose 11; pH 7.4 was adjusted with NaOH. Hippocampal slices were cut at a thickness of 200 μm by using a vibratome (VT1200S; Leica Biosystem, Muttenz, Switzerland). Whole-cell current-clamp and voltage-clamp recordings were made using a patch clamp amplifier (MultiClamp 700B; Molecular Devices, San Jose, CA, USA). Patch pipettes were filled with an intracellular solution of the following composition (in mM): K-gluconic acid 140, NaCl 10, MgCl_2_ 1, HEPES 10, EGTA 11, Mg-ATP 1.5, Li-GTP 0.3; pH 7.3 was adjusted with KOH. In the voltage-clamp recording mode of the amplifier, the holding potential of astrocytes was set to −80 mV, and that of neurons to −70 mV.

Astrocytes were discriminated from neurons by visual observation, and by their failure to fire action potentials in response to a supra-threshold depolarizing current injection. AMPA (100 μM), Bz-ATP (1000 μM) and muscimol (100 μM), all from Sigma-Aldrich, were applied locally to astrocytes and neurons by means of a computer-controlled solenoid valve-driven pressurized superfusion system (VC^3^8; ALA Scientific Instruments, Farmingdale, NY, USA). The drug application tip touched the surface of the brain slice and was placed 100-150 μm afar from the patched cell. Agonists were applied for 10 s every 3 min, in a low X^2+^ aCSF solution (MgCl_2_ was omitted from the medium and the CaCl_2_ concentration was decreased to 0.5 mM). When the same concentration of an agonist was applied twice, the mean current response was calculated for statistical evaluation.

### Recording of spontaneous postsynaptic currents (sPSCs) and spontaneous excitatory postsynaptic currents (sEPSCs)

sPSCs and sEPSCs were recorded from neurons at the holding potential of −70 mV as described previously [45]. They were analyzed by means of the pClamp 10.4 software package, by recording amplitudes exceeding the detection threshold set at three times the standard deviation above the baseline noise of the recordings. False positive noise-triggered fluctuations and signals with a non-monotonic rising phase and/or additional events within the decay phase were rejected on visual inspection (<2%). Since sPSCs compose of both action potential-induced and spontaneous vesicular glutamate/GABA release, the blockade of GABA_A_ receptors by gabazine (10 µM) left us with pure glutamatergic sEPSCs.

sPSCs/sEPSCs were recorded for 5 min in the absence and for another 5 min in the presence of Bz-ATP (300 µM). The mean of the pre-drug (control) sPSC/sEPSC amplitude and frequency was considered as 100% and was used as a reference value for calculating the percentage change caused by Bz-ATP. Once again, reversibility of the Bz-ATP effect was demonstrated by recording sPSCs/sEPSCs for another 5 min after washing out Bz-ATP.

### Materials

The drugs used were the following: JNJ-47965567, gabazine hydrobromide (Tocris Biosciences, Bristol, UK); kainic acid hydrate (MedChem Express, Monmouth Junction, NJ, USA); diazepam (Shanghai Xudong Haipu Pharmaceutical Co., Shanghai, China).

Kainic acid was dissolved in saline (0.9% NaCl) and JNJ-47965567 was dissolved in 30% sulfobutylether-β-cyclodextrin + 70% saline. Diazepam was supplied by the producer as a solution containing propylene glycol and ethanol as co-solvents.

### Data analysis

Data were analyzed using GraphPad Prism 10.0. Sample sizes (*n*) refer to the number of mice for GFAP and S100β quantification, or the number of individual astrocyte cells (derived from at least 3 mice per group) for morphological analyses. The normality of data distribution was assessed using the Shapiro-Wilk test. Two-groups were compared with the parametric Student’s *t*-test or the non-parametric Mann-Whitney test, as appropriate. Data with more than two groups were tested for significance using the one-way ANOVA test followed by the Tukey’s test. Multiple comparisons between data were tested in case of their non-normal distribution by the Kruskal-Wallis ANOVA on ranks, followed by the Dunn’s test. Data are expressed as mean ± SEM, and a P-value < 0.05 was considered statistically significant.

### Author Contributions

XL, MTK, PI, and YT planned the experiments and wrote the original version of the manuscript. SSL helped with the histological part of the study. ESV, BS, AV, and PR revised the manuscript. All authors read and approved the final version of the paper.

### Funding

We are grateful for financial support from the NSFC-RSF (82261138557), Sino-German Centre (GZ919, M-0679), the Sichuan Science and Technology Program (2025YFHZ0121, 2026NSFSC1857), and the China Postdoctoral Science Foundation (2024MD753906).

## Results

The hippocampus is preferentially damaged by KA-induced seizures in mice and is also preferentially afflicted by human temporal lobe epilepsy [46, 47]. We have used a dual approach in these experiments to characterize changes occurring in hippocampal astrocytes after KA-induced SE: On the one hand the astrocytes were filled with the endogenously synthesized fluorescent label mCherry (3 weeks after intra-hippocampal injection of the mCherry-carrying virus), and thereby we labelled also small astrocytic processes, which allows indirect conclusions on changes in leaflets by the use of laser scanning confocal microscopy and 3D reconstruction of the astrocytic profiles. Otherwise, the small leaflets are below the resolving power of the light microscopic observation of any fluorescent-stained specimen. On the other hand, we stained with conventional immunofluorescence the astrocytic preparations and identified them by their glial fibrillary acidic protein (GFAP) and glial-specific calcium-binding protein (S100β)-immunopositivity.

### Atrophy and down-regulation of ezrin expression in *stratum oriens* astrocytes after kanic acid injection, and the prevention of these effects by P2X7R blockade

Confocal microscopy was used not only for the visualization of mCherry positive puncta but also for the confirmation of ezrin-mCherry co-localization which is likely to be specific for leaflets (Fig. 2A). In the first series of experiments, we determined various astrocytic parameters 2 weeks after the i.p. injection of saline to the mice (NS, normal saline; KA, kainic acid; Fig. 2). Under these control conditions, we measured the astrocytic volume per µm^3^, and the length of the primary astrocytic branches from their somal origin or their bifurcation in µm (Fig. 2B, C). Similarly, we determined the extent of ezrin-immunoreactivity (IR) at the surface of the cell body of astrocytes (soma; Fig. 2D), and at the surface of their branches (Fig. 2E).

**Figure 2.**
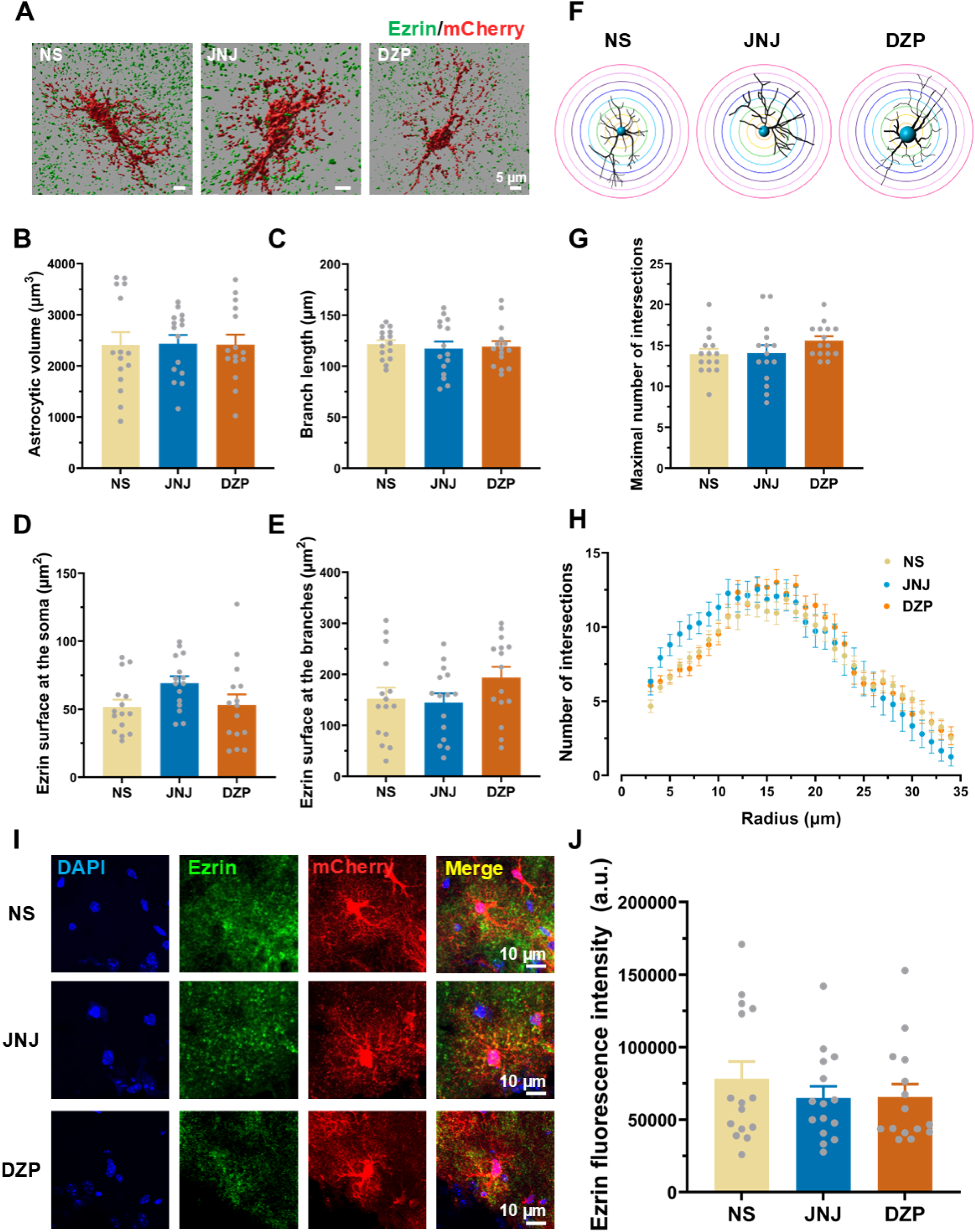
JNJ-47965567 and diazepam-treatment do not alter astrocyte morphology or ezrin immunofluorescence at astrocytes in the hippocampal *stratum oriens*. (A) Representative 3D surface reconstructions of hippocampal AAV-GfaABC1D-mCherry-labeled astrocytes (red) immunolabeled for ezrin (green) from mice treated with normal saline (NS), JNJ-47965567 (JNJ; 30 mg/kg, i.p.), or diazepam (DZP; 10 mg/kg, i.p.). Scale bar, 5 μm. (B-C) Morphometric quantification of astrocytes showing (B) astrocytic volume (P>0.05), (C) primary branch length (P>0.05). (D, E) Quantitative analysis of ezrin colocalization with astrocytes; (D) ezrin puncta area localized at the astrocytic soma (P>0.05); (E) ezrin puncta area localized to the astrocytic peripheral branches (P>0.05). (F) Schematic representation of Sholl analysis used to assess astrocytic complexity. (G) Maximal number of intersections (P>0.05. (B-E) One-way ANOVA followed by the Tukey’s test was used for statistical evaluation. (H) Intersection profile relative to the distance from the soma. (I) High-magnification confocal images showing ezrin (green) and mCherry-labeled astrocytes (red); DAPI-stained nuclei (blue). Scale bar, 10 μm. (J) Ezrin fluorescence intensity (expressed as area under the curve; a.u.) within the mCherry-defined astrocytic area (P>0.05; Kruskal-Wallis ANOVA followed by the Dunn’s test). All data are presented as mean±S.E.M. Dots represent individual astrocytic values (n=15).

The i.p. injection of the blood-brain permeable and highly specific P2X7R antagonist JNJ47965567 (30 mg/kg, i.p.; in the following JNJ), 1 h before the application of KA (30 mg/kg, i.p.) did not alter the above astrocytic parameters (Fig. 2B-E). JNJ is known to decrease systemic KA-induced SE under the present experimental conditions (see Fig. 1 in [30]). The benzodiazepine anti-SE drug diazepam (10 mg/kg, i.p.) also failed to modify the astrocytic volume, the branch length, as well as the surface staining by ezrin of the cell somata and branches (Fig. 2B-E).

Then we performed a Sholl analysis by laying at 1 µm distances concentric circles around the soma of each astrocyte and measured the number of intersections of these circles with the astrocytic branches (Fig. 2F). Once again JNJ and diazepam, both alone, failed to alter the number of intersections up to a radius of 35 µm in comparison with saline injection (Fig. 2H). For the maximum number of intersections this is documented in Fig. 2G. Eventually, the ezrin-mCherry fluorescent co-staining was determined in astrocytes, whose cell nuclei were labeled by the auto-fluorescent compound DAPI (Fig. 2I). The evaluation of ezrin fluorescence intensity did not show any change when the effects of JNJ or diazepam were compared with the values obtained under saline injection (Fig. 2J).

After having confirmed that neither JNJ nor diazepam modifies the mCherry labelling of astrocytes in the mouse hippocampus after saline injection, we turned our attention to possible long-term changes caused by SE, 2 weeks after KA injection, with and without the preceding application of JNJ or diazepam, both known to reduce/prevent epileptic fits. According to expectations, KA-induced SE led to profound astrocytic atrophy, as shown by the representative panels in Fig. 3A. The statistical evaluation of a number of experiments confirmed that the astrocytic volume, the branch length, and the maximal number of intersections as determined by a Sholl analysis, all decreased 2 weeks after KA injection when compared with the saline-injected controls (Fig. 3B-F).

**Figure 3.**
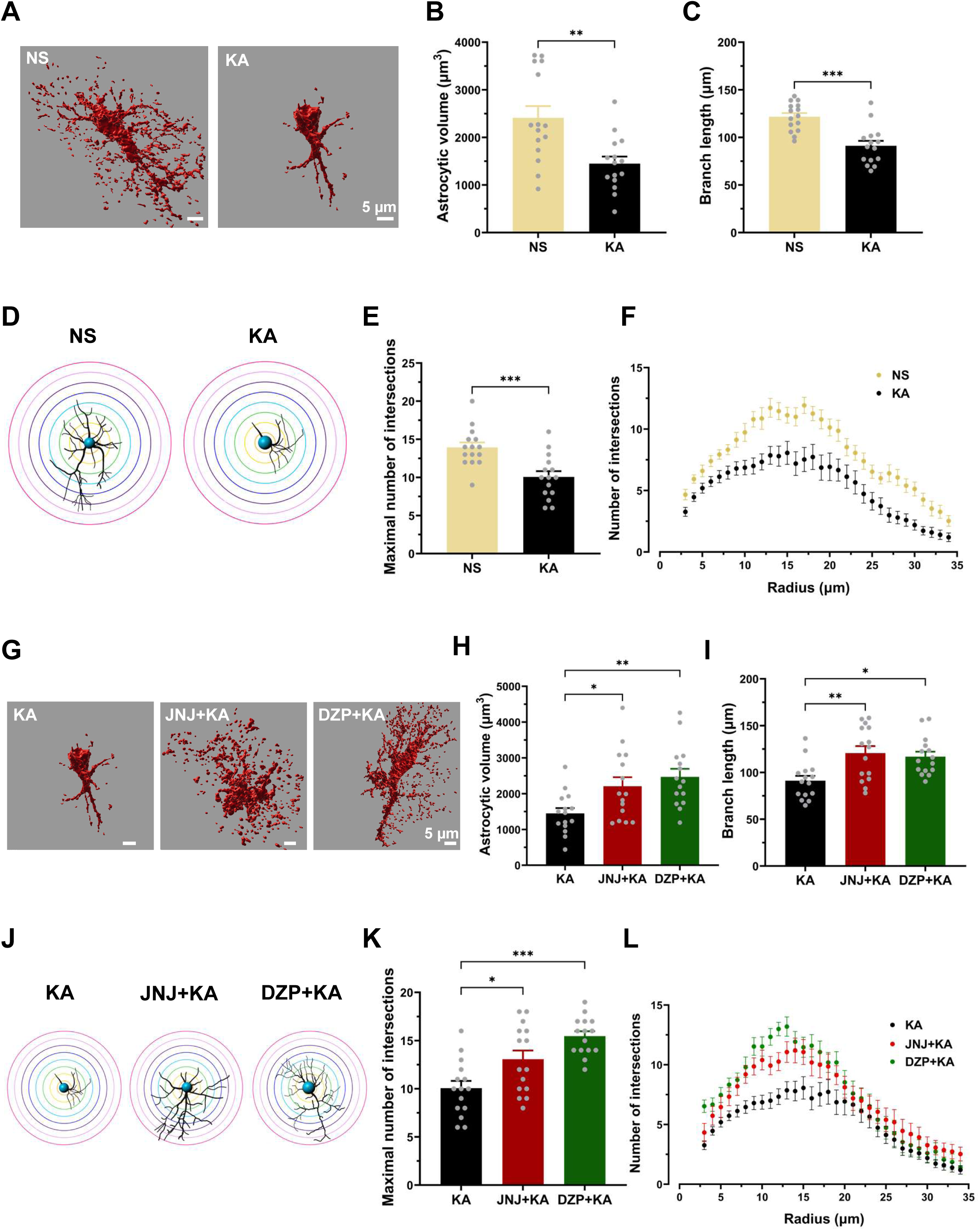
JNJ-47965567 and diazepam-treatment preserve astrocytic morphology after kainic acid-induced atrophy in the hippocampal *stratum oriens*. All abbreviations in this and the Figs. 4-5 were as defined in Fig. 2. (A) Representative 3D surface reconstructions of hippocampal mCherry-labeled astrocytes (red) from mice treated with NS or kainic acid KA. Scale bar, 5 μm. (B, C) Morphometric quantification of astrocytes showing (B) astrocytic volume (**P<0.01) and (C) primary branch length (***P<0.001). (D) Schematic representation of Sholl analysis used to assess astrocytic complexity. (E) Maximal number of intersections (***P<0.001). (B-E) The unpaired two-tailed t test was used for statistical evaluation. (F) Intersection profile relative to the distance from the soma. (G) Representative 3D surface reconstructions of hippocampal mCherry-labeled astrocytes (red) from KA-, JNJ+KA-, and DZP+KA-treated mice. Scale bar, 5 μm. (H, I) Morphometric quantification of astrocytes showing (H) astrocytic volume (*P<0.05; **P<0.01) and (I) primary branch length (*P<0.05, **P<0.01). (J) Schematic representation of Sholl analysis. (K) Maximal number of intersections (*P<0.05, ***P<0.001) (H-I, K) One-way ANOVA followed by the Tukey’s test was used for statistical evaluaton. (L) Intersection profile relative to the distance from the soma. All data are presented as mean±S.E.M. Dots represent individual astrocytic values (n=15).

At the same time the effect of KA disappeared when it was applied 1 h after a preceding injection of JNJ or diazepam (Fig. 3G-L). The findings with the selective P2X7R antagonist JNJ suggest that KA induced a massive release of ATP from damaged CNS cell types; this ATP acted at astrocytic P2X7Rs and caused their prominent atrophy. Similarly, in case of the pretreatment of mice with diazepam, which interfered with the effect of KA to cause seizures, there was no morphological injury of astrocytes caused by KA otherwise.

In the following experiments we investigated the co-expression of ezrin and mCherry in the 3D reconstructed puncta (Fig. 4A), and noticed that KA causes 2 weeks after its application, a marked decrease in the presence of ezrin puncta at the astrocytic branches (Fig. 4C), but not at the somata (Fig. 4B) of mCherry-labelled astrocytes, when compared with the effect of saline injection. The measurement of ezrin immunofluorescence-intensity also showed its marked decrease after KA application at the mCherry-positive cells (Fig. 4D, E).

**Figure 4.**
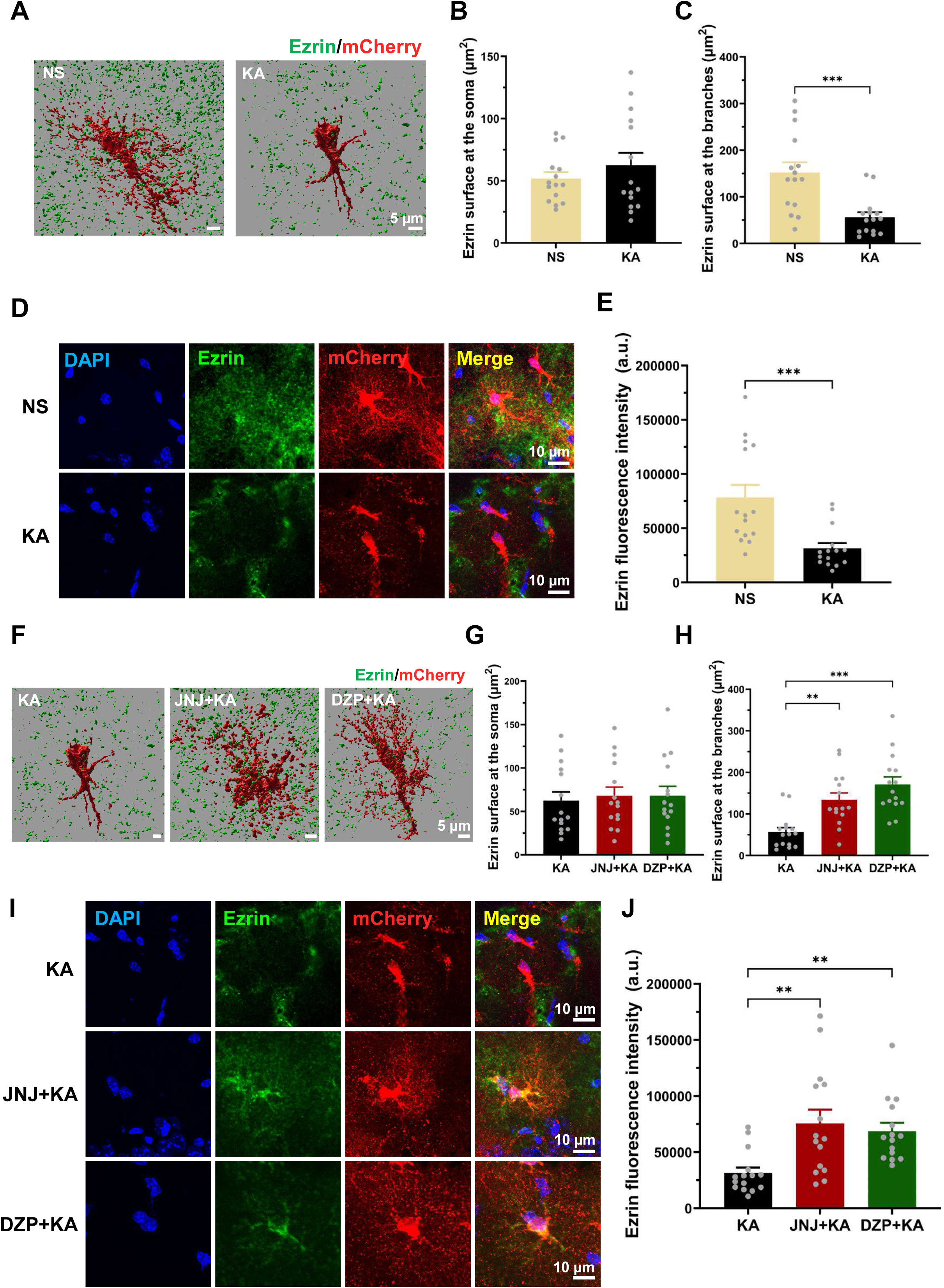
JNJ-47965567 and diazepam-treatment prevent the reduced expression of ezrin at astrocytic peripheral processes following kainic acid administration in the hippocampal *stratum oriens*. (A) Representative 3D surface reconstructions of hippocampal mCherry-labeled astrocytes (red) immunolabeled for ezrin (green) from mice treated with NS or KA. Scale bar, 5 μm. (B, C) Quantitative analysis of ezrin colocalization with astrocytes showing (B) ezrin puncta area localized to the astrocytic soma (P>0.05) and (C) ezrin puncta area localized to the astrocytic peripheral branches (***P<0.001). (D) Representative confocal images showing ezrin (green) and mCherry-labeled astrocytes (red); DAPI-stained nuclei (blue) in NS and KA groups. Scale bar, 10 μm. (E) Ezrin fluorescence intensity within the mCherry-defined astrocytic area (***P<0.001) (B, C, E) The Mann-Whitney test was used for statistical evaluation. (F) Representative 3D surface reconstructions of hippocampal mCherry-labeled astrocytes (red) immunolabeled for ezrin (green) from KA-, JNJ+KA-, and DZP+KA-treated mice. Scale bar, 5 μm. (G, H) Quantitative analysis of ezrin colocalization with astrocytes showing (G) ezrin puncta area at the soma (P>0.05) and (H) ezrin puncta area at the branches (**P<0.01, ***P<0.001). (I) Representative confocal images showing ezrin (green) and mCherry-labeled astrocytes (red); DAPI-stained nuclei (blue) in the KA, JNJ+KA, and DZP+KA groups. Scale bar, 10 μm. (J) Ezrin fluorescence intensity within the astrocytic area (**P<0.01) (G, H, J) The Kruskal-Wallis ANOVA followed by the Dunn’s test was used for statistical evaluation. All data are presented as mean±S.E.M. Dots represent individual astrocytic values (n=15).

In accordance with these findings, both JNJ and diazepam prevented the KA-induced decrease of ezrin expression, at the astrocytic branches (Fig. 4H), but had no effect at the somata of these cells (Fig. 4G). A representative picture in Fig. 4F convincingly documents these changes. Furthermore, Fig. 4I, J support these findings by yielding identical results by the quantitative evaluation of the ezrin immunofluorescence-data.

### Changes in GFAP- but not S100β-immunoreactivity in hippocampal astrocytes after kainic acid injection, and prevention of this effect by P2X7R blockade

Eventually, we investigated the changes in immunofluorescence intensity in *stratum oriens* and *stratum radiatum* astrocytes in the neighborhood of the hippocampal CA1 pyramidal cell region. We found that in the *stratum radiatum*, but not in the *stratum oriens* KA-injection increased the number of GFAP-positive astrocytes (Fig. 5A-C), while the number of S100β-positive astrocytes did not change in either region (Fig. 5D-F). This difference may be due to the fact that GFAP-IR marks astrocytes undergoing inflammatory reactions, whereas S100β is a more reliable marker of the general number of astrocytes [10]. Nonetheless, the selective increase of GFAP-positive astrocytic numbers by KA could be prevented by the pre-application of JNJ and thereby indicated the involvement of P2X7R activation in this process.

**Figure 5.**
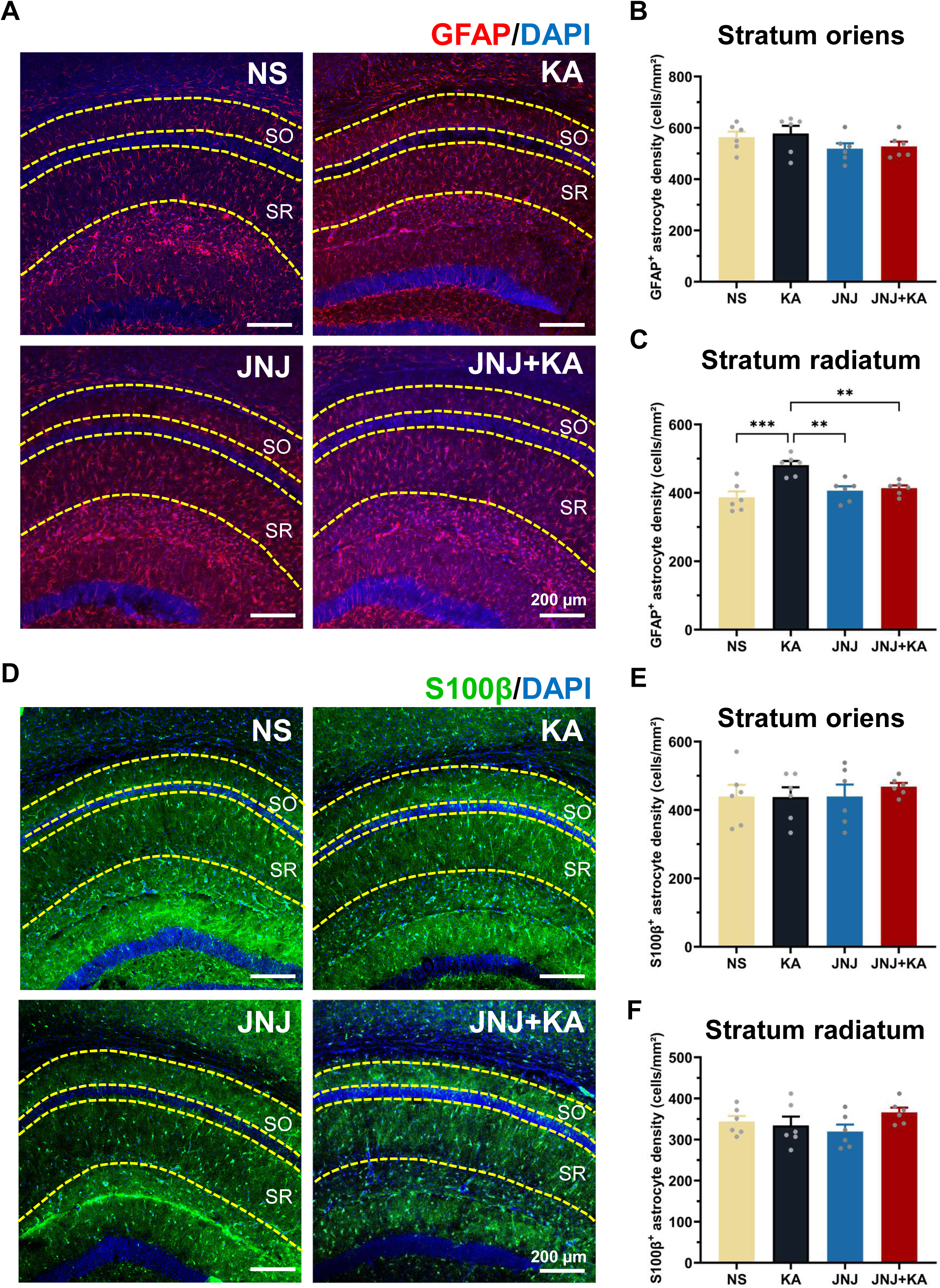
JNJ-47965567-treatment attenuates kainic acid-induced reactive astrogliosis in the hippocampal *stratum radiatum*, whereas there is no effect in the *stratum oriens*. (A) Representative immunofluorescence images of GFAP- (red) and DAPI-staining (blue) in the hippocampus from mice treated NS, KA, JNJ, or JNJ+KA. Dashed yellow lines delineate the *stratum oriens* (SO) and *stratum radiatum* (SR). Scale bar, 200 μm. (B, C) Quantitative analysis of GFAP-positive astrocytic density showing (B) in the SO (P>0.05) and (C) in the SR (**P<0.01, ***P<0.001; in both cases one-way ANOVA followed by the Tukey’s test). (D) Representative immunofluorescence images of S100β- (green) and DAPI-staining (blue) in the hippocampus from mice under the indicated treatment conditions. Dashed yellow lines indicate SO and SR. Scale bar, 200 μm. (E, F) Quantitative analysis of S100β-positive astrocytic density (E) in the SO (P>0.05) and (F) in the SR (P>0.05; in both cases one-way ANOVA followed by the Tukey’s test). All data are presented as mean±S.E.M. Dots represent individual values in 6 mice.

### Patch-clamp experiments demonstrate P2X7R-mediated potentiation of kainic acid-induced seizures in hippocampal *stratum oriens* astrocytes; no similar effect was observed in hippocampal CA1 neurons

In addition to measuring morphological and immunohistochemical changes in astrocytes after KA-induced SE, we also searched for functional alterations in this condition. For this purpose, we recorded by means of the whole-cell patch-clamp technique modifications in prototypic agonist-induced current amplitudes in hippocampal astrocytes and neurons. Diverse ionotropic receptors were stimulated by agonists, such as NMDA (NMDA-type glutamate-R; 100 µM), muscimol (GABA_A_-R; 100 µM) besides the stimulation of P2X7Rs by Bz-ATP (1000 µM). The brain slices were kept in a low X^2+^ aCSF, in order to increase the current amplitude evoked by Bz-ATP (Ren, 2023).

1 h after KA injection, the effect of Bz-ATP tended to increase in hippocampal astrocytes in comparison with the current responses to Bz-ATP in preparations taken from saline pre-treated mice but did not reach the level of statistical significance (Fig. 6A-C). However, there was a marked difference under these experimental conditions in preparations taken from mice 2 weeks after KA-injection, at a point in time when the morphometric-immunohistochemistry experiments were carried out (Fig. 6G). When the same procedure was repeated, but the brain slices were continuously superfused with the P2X7R antagonist A-438079 (10 µM), this difference disappeared. It was concluded that the Bz-ATP-induced currents became larger after KA-injection, due to the sensitization of astrocytic P2X7Rs by the damage-induced release of ATP.

**Figure 6.**
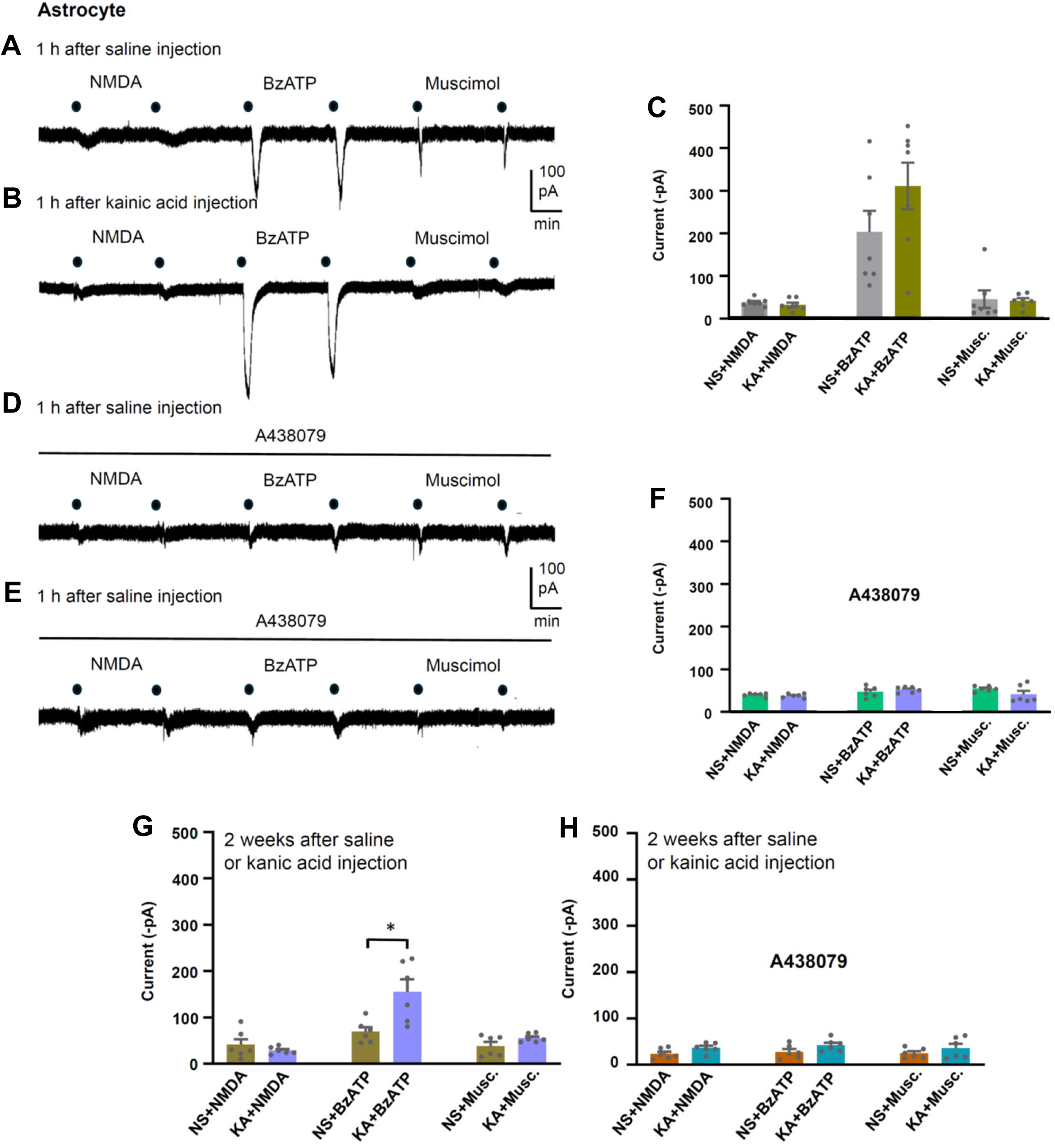
NMDA, Bz-ATP and muscimol-induced current responses in hippocampal *stratum oriens* astrocytes of mice. Each agonist was superfused twice for 10 s each with an inter-application interval of 3 min, at a holding potential of −80 mV. The concentrations of the agonists were 100 µM (NMDA, muscimol) and 1000 µM (Bz-ATP) in a low X^2+^ bath medium (see Methods). The current response was calculated for each agonist as a mean of the two subsequent applications. Mice were 1 h or 2 weeks before preparation of hippocampal brain slices injected with normal saline (NS) or kainic acid (KA, 30 mg/kg, i.p.). (A) Representative recording 1 h after saline injection. (B) Representative recording 1 h after KA injection. (C) Comparison of the effects of NMDA, Bz-ATP, and muscimol, 1 h after saline or KA-application. P>0.05 all, n=6; Kruskal-Wallis ANOVA, followed by the Dunn’s test. (D) Representative recording 1 h after saline injection in the continuous presence of A438079 (10 µM). (E) Representative recording 1 h after KA injection in the continuous presence of A438079 (10 µM). (F) Comparison of the effects of NMDA, Bz-ATP, and muscimol, 1 h after saline or KA-application in the continuous presence of A438079 (10 µM). P>0.05 all, n=6; one-way ANOVA followed by the Tukey’s test. (G) Comparison of the effects of NMDA, Bz-ATP, and muscimol, 1 h after saline or KA-application. *P<0.05, as indicated; n=6; Kruskal-Wallis ANOVA, followed by the Dunn’s test. (H) Comparison of the effects of NMDA, Bz-ATP, and muscimol, 1 h after saline or KA-application in the continuous presence of A438079 (10 µM). P>0.05 all, n=6; one-way ANOVA followed by the Tukey’s test. For further details see the Materials and Methods Section. All data are presented as mean±S.E.M. Dots represent individual values in 6 mice.

**Figure 7.**
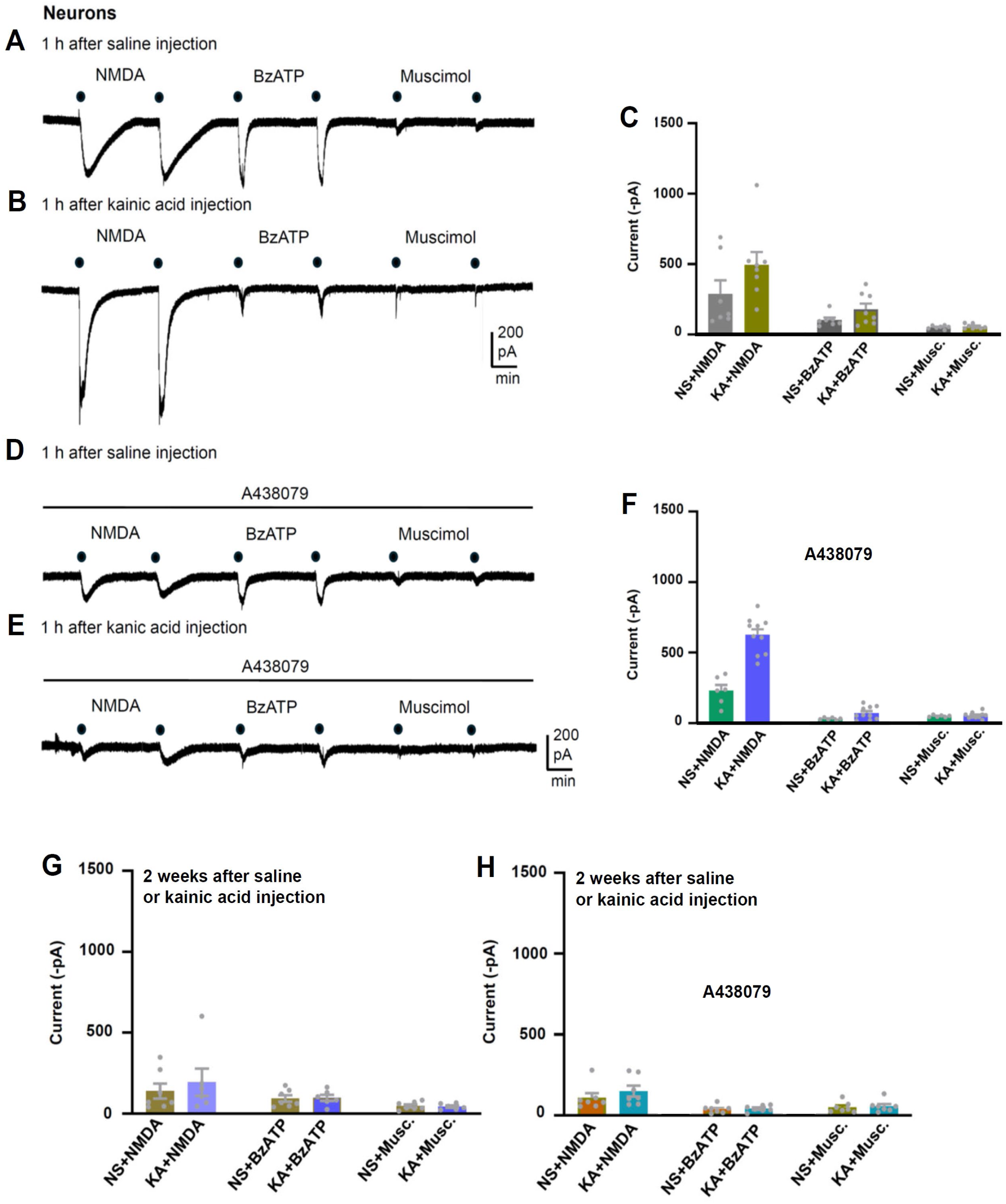
NMDA, Bz-ATP and muscimol-induced current responses in hippocampal CA1 pyramidal neurons of mice. For methodological details see the Legend to Fig. 6 and the Materials and Methods Section. (A) Representative recording 1 h after saline injection. (B) Representative recording 1 h after KA injection. (C) Comparison of the effects of NMDA, Bz-ATP, and muscimol, 1 h after saline or KA-application. P>0.05 all, n=6; Kruskal-Wallis ANOVA, followed by the Dunn’s test. (D) Representative recording 1 h after saline injection in the continuous presence of A438079 (10 µM). (E) Representative recording 1 h after KA injection in the continuous presence of A438079 (10 µM). (F) Comparison of the effects of NMDA, Bz-ATP, and muscimol, 1 h after saline or KA-application in the continuous presence of A438079 (10 µM). ***P>0.001, as indicated, n=6-10; Kruskal-Wallis ANOVA, followed by the Dunn’s test. (G) Comparison of the effects of NMDA, Bz-ATP, and muscimol, 1 h after saline or KA-application. P>0.05 all; n=6; one-way ANOVA followed by the Tukey’s test. (H) Comparison of the effects of NMDA, Bz-ATP, and muscimol, 1 h after saline and KA-application in the continuous presence of A438079 (10 µM). P>0.05 all, n=6; one-way ANOVA followed by the Tukey’s test. All data are presented as mean±S.E.M.

In contrast to astrocytes, neurons responded to NMDA in the continuous presence of the P2X7R antagonistic A438079 in brain slices taken from KA-treated mice, with larger current responses than in preparations taken from saline-treated animals. It appears that the strong reduction of the Bz-ATP-current unmasked in neurons, but not in astrocytes, a marked potentiation of the NMDA-current responses. There was no similar difference observed in the neurons of brain slices taken from mice injected 2 weeks before with saline or KA.

### P2X7R-activation increases the sPSC/sEPSC current but not amplitude in hippocampal CA1 neurons

In the last series of experiments, we recorded sPSCs (caused by the spontaneous release of glutamate and GABA) from hippocampal neurons kept in brain slices. Neither the amplitude nor the frequency of these events differed from their pre-drug values, during superfusion with Bz-ATP (300 µM) for 5 min (Fig. 8A-C). However, when we calculated the percentage change of the amplitude and frequency of the sPSCs, it became evident that Bz-ATP increased the sPSC frequency but not amplitude in the saline, as well as the KA-treated (both 1h and 2 weeks after KA injection) preparations Fig. 8D). Nonetheless, this potentiation was the same under all three experimental conditions.

**Figure 8.**
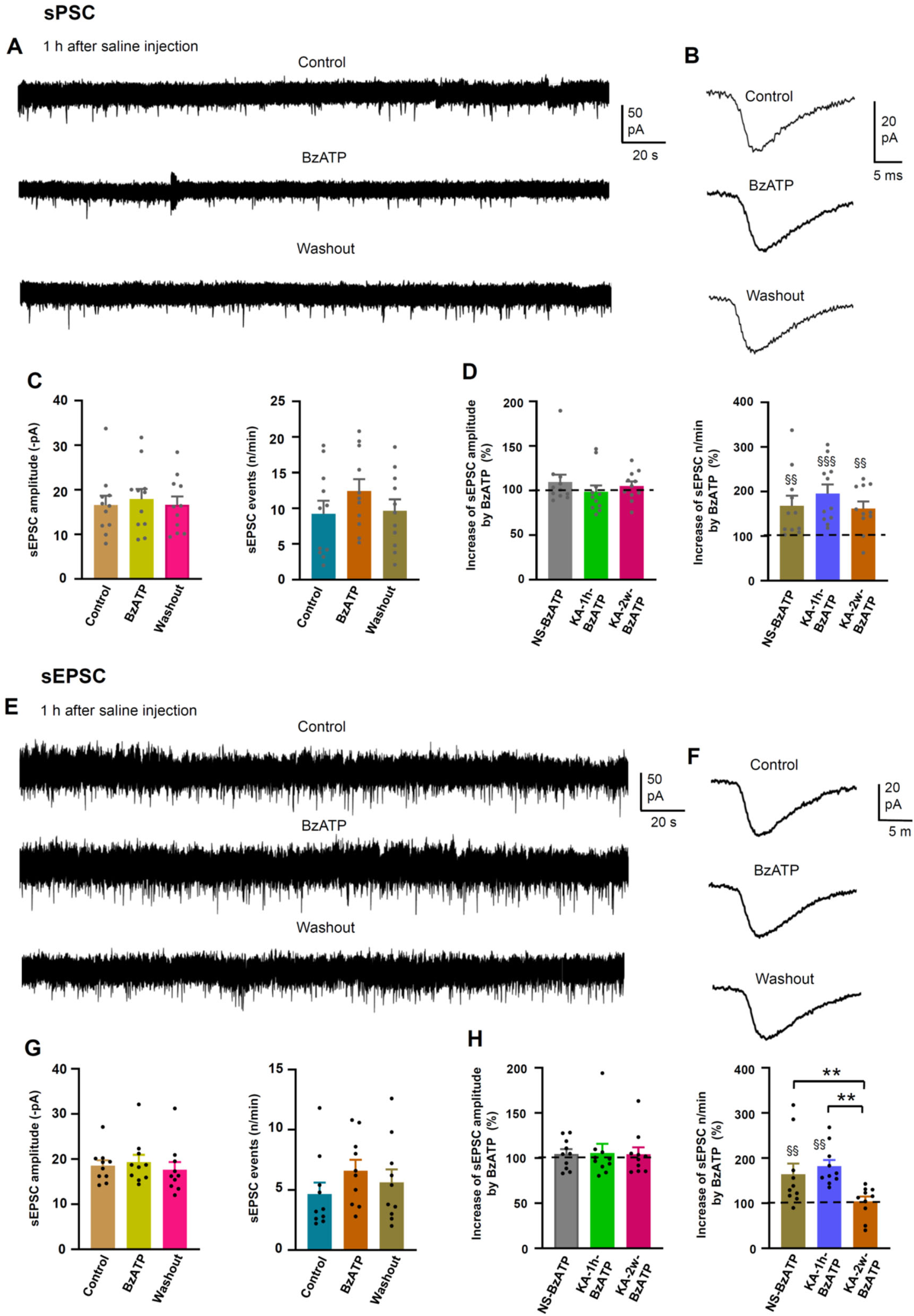
Spontaneous postsynaptic currents (sPSCs) and spontaneous excitatory postsynaptic currents (sEPSCs) of hippocampal CA1 pyramidal neurons; effects of Bz-ATP. sEPSCs were recorded in the continuous presence of gabazine (10 µM). Bz-ATP (300 µM) effects on the amplitude and frequency of the spontaneous currents in low X^2+^ aCSF. (A) Representative recordings or sPSCs before, during, and after superfusion with Bz-ATP for 5 min, as well as after washing out this agonist. (B) Representative, averaged sPSC amplitude before, during and after a 5 min superfusion with Bz-ATP. (C) Mean±S.E.M. sPSC amplitudes (left panel) and mean±S.E.M. frequencies (right panel) calculated from 11 cells. P>0.05 all; one-way ANOVA, followed by the Tukey’s test in both panels. (D) Percentage changes of sPSC amplitudes (left panel; P>0.05 all, Kruskal-Wallis ANOVA, followed by the Dunn’s test) and frequency (right panel; P>0.05 all; one-way ANOVA, followed by the Tukey’s test; ^§§^P<0.01; Kruskal-Wallis ANOVA, followed by the Dunn’s test), as calculated relative to the pre-Bz-ATP amplitude and frequency set at 100%, and shown as mean±S.E.M. values of 11 cells. (E) Representative recordings or sEPSCs before, during, and after superfusion with Bz-ATP for 5 min, as well as after washing out this agonist. (F) Representative, averaged sEPSC amplitude before, during and after a 5 min superfusion with Bz-ATP. (G) Mean±S.E.M. sEPSC amplitudes (left panel) and mean±S.E.M. sEPSC frequencies (right panel) calculated from 11 cells. P>0.05 all; one-way ANOVA, followed by the Tukey’s test in both panels. (H) Percentage changes of sEPSC amplitudes (left panel; P>0.05 all, Kruskal-Wallis ANOVA, followed by the Dunn’s test) and frequency (right panel; P>0.05 all; one-way ANOVA, followed by the Tukey’s test; ^§§^P<0.01; Kruskal-Wallis ANOVA, followed by the Dunn’s test) were calculated relative to the pre-Bz-ATP amplitude and frequency set at 100%, and shown as mean±S.E.M. values of 11 cells.

When the GABAergic component of the sPSCs was blocked by the continuous presence of gabazine (10 µM; Fig. 8E-G), the resulting sEPSC amplitudes and frequencies reacted in a similar manner to Bz-ATP than the sPCs. The only difference was that 2 weeks after KA-application, the Bz-ATP-induced percentage increase in frequency of sPSCs returned to its pre-drug level after the washout of Bz-ATP (Fig. 8H). In view of the absence of a change in sPSC or sEPSC amplitudes by Bz-ATP both in saline- and KA-treated mice, but the pronounced facilitation of the sPSC/sEPSC frequency by Bz-ATP in both cases, we concluded that P2X7R-stimulation might have a pre-synaptic facilitatory effect on glutamate and probably also GABA release, which was not altered by KA pre-treatment of mice. Although Bz-ATP is not a selective P2X7R agonist, its potentiation of the sPSC/sEPSC frequency was due to the stimulation of P2X7Rs, as confirmed by the abolition of this effect by A-438079 [45].

## Discussion

The main purpose of this paper was to clarify, whether the KA-induced atrophy of astrocytes probably by preferential damage to the PAPs and therefore to the synaptic cradle is the immediate reason for excitability changes in hippocampal CA1 neurons (see [10, 11]. An important driver of these events might be the massive release of ATP in the hippocampus which *via* stimulation of astrocytic P2X7Rs might trigger reverberating and self-strengthening neuronal circuits, resulting in epileptic discharges.

P2X7Rs have the lowest affinity for ATP among all P2X and P2YRs, a feature that defines it as a pathological receptor activated only by large local concentrations of ATP, rising to levels at sites of inflammation, cancer, and in fact SE, to heights needed for receptor-stimulation [48–51]. The P2X7R is a ligand-gated cationic channel which allows the passage of small cations Na^+^, K^+^, and Ca^2+^ [48], but allows also the passage of large cationic molecules immediately from its initial activation, but at a slower pace than that of the smaller molecules [52, 53]. This property of the P2X7R is getting prominent during repetitive or long-lasting contact with its agonist ATP. The P2X7R causes necrosis/apoptosis and neuroinflammation, the latter mostly because of interleukin-1β (IL-1β) secretion; the gating of the P2X7R-channel by extracellular ATP allows the efflux of large amounts of intracellular K^+^ that in turn drives the assembly of the NLRP3 inflammasome and the activation of caspase-1 [54, 55]. Caspase-1 catalyzes the conversion of pro-IL-1β to the active metabolite IL1β; the stimulus for pro-IL-1β synthesis is the occupation of the toll-like receptor-4 (TLR-4) by lipopolysaccharide. Therefore, IL-1β secretion is based on the two-hit concept, requiring the simultaneous activation of TLR-4 and P2X7Rs. P2X7Rs amplify CNS damage in neurodegenerative diseases, because of their necrotic (loss of intracellular ions and metabolites), apoptotic (activation of the caspase intracellular cascade), and neuroinflammatory (release of IL1β and IL-6) function superimposed on the basic production of toxic metabolites e.g. in Alzheimer’s disease (amyloid-β), Parkinson’s disease (α-synuclein), etc. [53, 56].

P2X7R-IR has been shown to be increased in hippocampus and cortex after experimentally induced SE [24, 57, 58]. However, the informative value of these experiments is rather limited in view of the questionably selective labeling of P2X7Rs by polyclonal antibodies used for immunohistochemistry investigations [59]. For some time, it was hoped that the use of transgenic P2X7R-enhanced green fluorescence protein (EGFP)-expressing neurons after chemically-induced SE will furnish strong evince for the up-regulation of P2X7Rs. Although the use of a first strain of transgenic mice appeared to support this assumption [60, 61], the generation of a second strain of transgenic mice raised serious doubts on the reliability of these findings [62, 63]. These dissonant results already highlight a major dispute on the existence of P2X7Rs in neurons in addition to their confirmed presence in microglia and astrocytes. While a group of authors favored the neuronal expression of P2X7Rs [64] another group refuted their arguments [65]. It was suggested that astrocytic P2X7Rs might release glutamate/GABA, stimulating their own receptors situated at neighboring astrocytes e.g. in the CA1 region of the hippocampus or the *substantia gelatinosa* of the spinal cord, instead of direct stimulation of neuronal P2X7Rs by their agonist ATP [45, 66]. Apparently, this discussion is not yet brought to an end, because after the intra-amygdala or i.p. injection of KA, P2X7Rs were identified both at hippocampal interneurons and microglial cells, the former exerting protective, while the latter pro-inflammatory effects [29].

The present results are in accordance with the assumption, although do not prove it unequivocally, that astrocytic P2X7Rs are the primary target of SE-induced release of ATP. The detailed morphometric analysis of the 3D data generated by means of mCherry expression in hippocampal astrocytes showed a decrease in astrocytic volume, branch length, and by Sholl analysis, a decrease of the maximal number of process intersections with concentric circles drawn around the cell soma, after KA application. All these changes were prevented by the injection of the anti-seizure drug diazepam or the P2X7R antagonist A438079 to mice preceding KA application. In addition, evaluation of the co-expression of ezrin (a plasmalemmal-cytoskeleton linker in astrocytes; [12, 13]) with mCherry led to identical conclusions. Thus, KA-SE caused astrocytic atrophy and suggested damage to PAPs *via* the release of ATP from injured CNS cells.

The recording of Bz-ATP-induced current responses in astrocytes and neurons also support this hypothesis. Comparison of Bz-ATP-current amplitudes in hippocampal brain slices prepared from saline- and KA-treated mice, 2 weeks after their application, indicated a potentiation by KA-induced seizures. The P2X7R antagonist A-438079 abolished the difference between astrocytic Bz-ATP-currents at this point in time. In contrast to the results in astrocytes, the Bz-ATP-currents measured both 1h and 2 weeks after KA-injection failed to differ in a statistically significant manner. Further, the facilitation of sPSC and sEPSC frequencies by KA-SE indicated a pre-synaptic potentiation of the spontaneous release of glutamate. This effect could be the consequence of an astrocytic modulatory effect and accompany the damage to astrocytes and the ensuing disturbance of neuronal homeostasis.

Astrocytes regulate synaptogenesis, support synaptic transmission through numerous transporters that control the local concentration of neuro(glio)transmitters in the synaptic cleft, provide ionostasis, supply neurotransmitter precursors and energy substrates, and regulate synaptic extinction [67–69]. Astrocytic atrophy is manifest in various neuropsychiatric and neurodegenerative disorders; the failure of astrocytic homeostatic support is the leading reason for neuronal damage and death in many CNS diseases, including SE.

There are multiple mechanisms by which astrocytes might increase neuronal excitability and drive epilepsy: These are in the first line (1) the astrocytic release of glutamate which possibly synchronizes neuronal activity, (2) the disturbance of a functionally linked water and potassium homeostasis, and (3) the uncoupling of astrocytes by the loss of gap-junction connections in the astrocytic syncytium [70–72]. ATP *via* the activation of P2X7Rs may induce inflammation in the CNS and cause reactive astrogliosis critically involved in the establishment of an epileptic focus [71, 73]. P2X7R antagonism thereby may act in an anti-epileptic manner primarily because of its anti-inflammatory action especially prominently in astrocytic networks.

In conclusion, our morphological, immunohistochemical and electrophysiological experiments strongly suggest that the establishment of SE in mice critically relies on the activation of astrocytic P2X7Rs in an epilepsy-relevant area of the brain, the hippocampus. In consequence, the blockade of this type of purinergic receptor is a promising alternative target, also in human therapy, to classic anti-epileptic drugs, which act only in a subgroup of patients and carry the danger of considerable side effects [74, 75].

## Availability of Data and Materials

All data can be obtained from the senior authors on reasonable request.

